# Widespread occurrence of bovine-like and new viruses in wild deer across the United States

**DOI:** 10.64898/2026.07.17.739239

**Authors:** Axel O.G. Hoarau, Meggan E. Craft, Simona J. Kraberger, Brock Geary, Jennifer Høy-Petersen, Julie C. Ellis, Jeremy Alder, Sara Hathaway, George Wittemyer, Guillaume Bastille-Rousseau, Tadao Kishimoto, Taylor Anderson, Travis Gallo, Jennifer M. Mullinax, Amira A. Roess, James D. Forester, Tyler J. Garwood, Tiffany M. Wolf, Jennifer L. Malmberg, Marnee K. Roundtree, Georgia C. Titcomb, Rebecca M. Windell, Maria A. Diuk-Wasser, Laura D. Plimpton, Meredith C. VanAcker, Alec Baker, Avery M. Corondi, W. David Walter, Tyler S. Walters, Daniel M. Grove, Justin R. Kosiewska, Dailee L. Metts, Cameron Mitchell, Lisa I. Muller, Mark Q. Wilber, Jacob D. Wyrick, Kezia R. Manlove, Megan Morrison, Lauren Smith, Virginia Stout, Grete Wilson-Henjum, Kim M. Pepin, Roderick B. Gagne

**Affiliations:** Department of Pathobiology, Wildlife Futures Program, University of Pennsylvania School of Veterinary Medicine, New Bolton Center, Kennett Square, PA, 19348, USA; Department of Ecology, Evolution and Behavior, University of Minnesota, Saint-Paul, MN, 55108, USA; The Biodesign Center for Fundamental and Applied Microbiomics, School of Life Sciences, Arizona State University, Tempe, AZ, 85281, USA; Department of Fish, Wildlife and Conservation Biology, Colorado State University, Fort Collins, CO, 80523, USA; Center for Wildlife Sustainability Research, Southern Illinois University, Carbondale, IL, 62901, USA; Department of Geography and Geoinformation Science, George Mason University, Fairfax, VA, 22030, USA; Department of Environmental Science and Technology, University of Maryland, College Park, MD, 20742, USA; Department of Global and Community Health, College of Health and Human Services, George Mason University, Fairfax, VA, 22030, USA; Department of Fisheries, Wildlife and Conservation Biology, University of Minnesota, Saint-Paul, MN, 55108, USA; Department of Veterinary Population Medicine, College of Veterinary Medicine, University of Minnesota, Saint-Paul, MN, 55108, USA; United States Department of Agriculture, Animal and Plant Health Inspection Services, Wildlife Services, National Wildlife Research Center, Fort Collins, CO, 80521, USA; Nebraska Game and Parks Commission, Lincoln, NE, 68522, USA; Department of Ecology, Evolution, and Environmental Biology, Columbia University, New York, NY, 10027, USA; Department of Evolution, Ecology, and Organismal Biology, University of California, Riverside, CA, 92521, USA; Pennsylvania Cooperative Fish and Wildlife Research Unit, The Pennsylvania State University, University Park, PA, 16802, USA; Dark Hollow Wildlife Consulting, Auburn, PA, 17922, USA; Department of Ecosystem Science and Management, The Pennsylvania State University, University Park, PA 16802; School of Natural Resources, University of Tennessee Institute of Agriculture, Knoxville, TN, 37996, USA; Department of Wildland Resources and Ecology Center, Utah State University, Logan, UT, 84322, USA; Utah Division of Wildlife Resources, Salt Lake City, UT, 84103, USA

**Keywords:** Wildlife, Virome, Metagenomic, *Odocoileus virginianus*, *Odocoileus hemionus*, Deer, Virus

## Abstract

Over the past several decades, deer populations in North America have grown considerably, resulting in frequent contact with humans and livestock, and increased potential for pathogen spillover. Despite the importance of pathogen spillover among humans, wildlife, and livestock, the diversity of viruses present in deer remains largely unknown. Using a metagenomic high-throughput sequencing approach, we characterized viral communities in the upper respiratory tracts of live mule deer (*Odocoileus hemionus*) and white-tailed deer (*Odocoileus virginianus*) captured at fourteen study sites in nine states across the United States. We identified vertebrate-infecting viruses in both deer species at all but two of the study sites, located in Illinois and Utah. Viral richness did not vary among species or study sites. However, viral community composition was different across study sites but not among deer species. Amongst the detected viral sequences, several originate from or were closely related to viruses previously described in humans (*e.g.*, severe acute respiratory syndrome coronavirus 2) and livestock (*e.g.*, bovine-like coronavirus). We also documented viruses recently discovered in deer, such as CHeRI orbivirus 1. Finally, we identified several new putative viruses in the *Picornaviridae*, *Rhabdoviridae* and *Tobaniviridae* families, including a novel *Aphthovirus* related to bovine rhinitis A virus in both deer species, across seven states and nine study sites. Our findings expand the understanding of viral diversity in two deer species across the United States, providing important insights for managing pathogens at the wildlife, livestock, and human interface.

## Introduction

Understanding of viral biodiversity has grown as metagenomic sequencing permits non-targeted detection across diverse hosts and environments (1–4). Yet only a small fraction of viruses has been described. Current knowledge of animal viromes remains limited to a small number of taxa, and most studies are primarily descriptive (5). As a result, it is challenging to draw comprehensive conclusions about their ecology and evolution (2). Comprehensive cataloging of all viruses, however, is not feasible. Instead, greater emphasis on focused, hypothesis-driven investigation of the ecological (*e.g.*, habitat use, host density) and evolutionary (*e.g*., host susceptibility, viral host plasticity) processes shaping virome composition in nature is needed to yield more generalizable insights (6). Such approaches can improve our understanding of viral emergence and support more targeted surveillance efforts. Given the threat viruses pose to the health of wild and domestic animals, humans, and ecosystems, a more thorough understanding of which viruses circulate in animal populations is important for contextualizing zoonotic risk and informing surveillance strategies.

Studying viromes in species with high population densities, broad habitat use, and frequent interactions with agricultural animals and humans, provides opportunities to identify viruses of potential concern (7, 8). In North America, deer (family Cervidae) occur at high abundance and occupy extensive geographic ranges (9, 10), in part due to intensive management efforts including predator control and regulated hunting (11). High population densities paired with extensive overlap with humans and livestock increase the risk of pathogen spillover (12–14). For example, human–deer spillover was documented during the COVID-19 pandemic. Severe acute respiratory syndrome coronavirus 2 (SARS-CoV-2) spilled over from humans into free-ranging deer on several occasions, with documented evidence of deer–to–deer and deer–to–human transmissions (15–18). Shortly after the start of the pandemic, the virus was widely detected in deer across the United Sates of America (USA) (19, 20). In other cases, lethal pathogens, such as *Mycobacterium bovis*, have spilled over between livestock and deer, resulting in mortality in one or other of these species (21–23). Despite the increased awareness of pathogen transmission at the deer–human–livestock interface, the deer virome remains largely unknown. Characterizing viral communities in deer could identify viruses shared among deer, humans and domestic animals, and elucidate the potential for viral recombination and emergence of novel viruses.

Characterizing deer viromes not only has important implications for the health of livestock, humans, and other wildlife, but could also provide valuable insight into environmental and evolutionary drivers shaping animal viromes (6). Deer live in diverse habitats ranging from large cities to vast wilderness areas, spanning a wide range of key biotic and abiotic conditions that may influence the diversity and structure of viral communities (6). Focusing on closely related but distinct deer species also provide an opportunity to investigate evolutionary constraints on virome composition (6, 24). Areas where multiple deer species occur sympatrically, could help researchers comparing the influence of geography and ecology with that of the host species in shaping virome structure. To explore these determinants, we leveraged a unique sampling design and a metagenomic high-throughput sequencing approach, to characterize viral communities in the upper respiratory tracts of mule deer (MuD; *Odocoileus hemionus*) and white-tailed deer (WTD; *Odocoileus virginianus*) captured in various environments and habitats across the USA.

## Material and Methods

### Sample collection

The samples used in this study were collected as part of a national longitudinal study entitled “Coordinated, targeted surveillance of pathogens in wild deer populations”, conducted throughout the USA (https://www.targetedsurveillance.com/; (25)). Details regarding the study design, deer capture, and sample collection protocols are available in Pepin *et al*., 2025 (25). We focused on deer captured during the winter of 2023–2024 (December to April) across nine states and fourteen study sites (**Table 1, Supplementary Table 1**). These sites represent diverse geographic and ecological contexts, including rural *versus* urban deer populations and varying degrees of interaction with humans and domestic animals (25). We restricted samples to those collected from live animals as conducting virome studies on dead animals may create biases towards individuals that were diseased, and viral composition may vary meaningfully after death (26). Nasopharyngeal swabs were collected from MuD and WTD using Fisherbrand™ (Fisher Scientific) or similar polyester-tipped applicators. Swabs were placed in 2 mL cryogenic vials (Cornig) containing 1.5 mL of DNA/RNA shield (Zymo Research). In accordance with manufacturer’s recommendations, samples were stored at room temperature for up to three weeks or frozen at −20°C before being shipped to the Wildlife Futures Program at New Bolton Center, Kennett Square, Pennsylvania, USA. Upon arrival, all samples were placed in a −80°C freezer until processed.

**Table 1:** Details on state, collection site, sampled species and number of samples included in each library/pool.

| Pool/Library ID | State | Collection site | Species | Number of samples I |
| --- | --- | --- | --- | --- |
| CO1 | Colorado | Poudre River | Mule deer (*Odocoileus hemionus*) | 10 |
| CO2 | Colorado | Poudre River | White-tailed deer (*Odocoileus virginianus*) | 10 |
| IL1 | Illinois | Shelbyville | White-tailed deer (*Odocoileus virginianus*) | 10 |
| IL2 | Illinois | Touch of Nature | White-tailed deer (*Odocoileus virginianus*) | 10 |
| IL3 | Illinois | Touch of Nature | White-tailed deer (*Odocoileus virginianus*) | 10 |
| IL4 | Illinois | Touch of Nature | White-tailed deer (*Odocoileus virginianus*) | 10 |
| MD | Maryland | DC area | White-tailed deer (*Odocoileus virginianus*) | 10 |
| MN1 | Minnesota | Carver Park | White-tailed deer (*Odocoileus virginianus*) | 10 |
| MN2 | Minnesota | Shakopee | White-tailed deer (*Odocoileus virginianus*) | 10 |
| NE | Nebraska | Carter Canyon | Mule deer (*Odocoileus hemionus*) | 10 |
| NY | New York | Staten Island | White-tailed deer (*Odocoileus virginianus*) | 10 |
| PA1 | Pennsylvania | Bedford/Fulton | White-tailed deer (*Odocoileus virginianus*) | 10 |
| PA2 | Pennsylvania | Treasure Lake | White-tailed deer (*Odocoileus virginianus*) | 10 |
| PA3 | Pennsylvania | Bedford/Fulton | White-tailed deer (*Odocoileus virginianus*) | 10 |
| PA4 | Pennsylvania | Bedford/Fulton | White-tailed deer (*Odocoileus virginianus*) | 10 |
| TN | Tennessee | Ames Plantation | White-tailed deer (*Odocoileus virginianus*) | 10 |
| UT1 | Utah | Pine Valley | Mule deer (*Odocoileus hemionus*) | 10 |
| UT2 | Utah | Pine Valley | Mule deer (*Odocoileus hemionus*) | 10 |
| UT3 | Utah | Pine Valley | Mule deer (*Odocoileus hemionus*) | 10 |
| UT4 | Utah | La Sals | Mule deer (*Odocoileus hemionus*) | 10 |
| UT5 | Utah | Nebo | Mule deer (*Odocoileus hemionus*) | 10 |
| **Total** |  |  |  | **210** |

### RNA extraction

We pooled samples by state, study site, and deer species. We randomly selected 10 samples per pool and, where possible, balanced the sex ratio within pools. We assigned one or three pools per study site, depending on the number of available samples (**Table 1, Supplementary Table 1**). For Poudre River, Colorado, the only study site where samples were available from sympatric MuD and WTD populations, we assigned one pool to each species.

We thawed samples at 4°C and used 140 µL of supernatant to extract RNA using the QIAamp viral RNA Mini Kit (QIAGEN). We eluted the RNA in 2 × 40 µL of AVE buffer to increase RNA yield. We quantified the extracted RNA using the Qubit RNA High Sensitivity Assay kit (Thermo Fisher Scientific) on a Qubit 4 Fluorometer (Thermo Fisher Scientific) (**Supplementary Table 1**). As our main objective was to prioritize viral detection over quantitative comparisons, we pooled RNA from each sample by equal volume (10 µL) into a single tube according to the pooling plan. We gently mixed the RNA by pipetting up and down and checked the final RNA concentration in each pool (**Supplementary Table 1**).

### Library preparation and sequencing

We treated each RNA pool with Ambion DNase I RNase-free (Thermo Fisher Scientific) to digest high molecular weight genomic DNA. We scaled the reaction to contain 1X DNase I buffer and 2 Units (U) of DNase I per 100 µL. We incubated the reactions at 37°C for 5 min, as recommended in previous studies (27), then immediately cleaned the mixture with Agencourt RNAClean XP beads (Beckman Coulter) using freshly prepared 80% ethanol. We eluted the samples in RNase-free water and used 10 µL of each pool for library preparation using the Illumina Stranded Total RNA Prep Ligation with Ribo-Zero Plus Microbiome (Illumina) combined with IDT for Illumina DNA/RNA UD Indexes (Illumina). We performed the incubation steps and PCR amplifications on a ProFlex 3 × 32-well thermocycler (Thermo Fisher Scientific). We assessed library quality using D1000 ScreenTapes (Agilent) on a 4150 TapeStation (Agilent). We measured the DNA yield of each library using the Qubit 1X dsDNA Assay kit (Thermo Fisher Scientific) on a Qubit 4 Fluorometer (Thermo Fisher Scientific) and stored them individually at −20°C. We then sent them to the Center for Host-Microbial Interactions, University of Pennsylvania, Philadelphia, Pennsylvania, USA, for sequencing. The libraries were pooled equimolarly and sequenced with paired-end sequencing (150 bp) on a NextSeq 2000 sequencing platform (Illumina).

### Bioinformatic pipeline and viral identification

We discarded low quality reads and trimmed adapters using fastp v0.24.0 (28). We then used Kneaddata v0.12.2 (https://bitbucket.org/biobakery/kneaddata) to remove rRNA and host reads, using the SILVA database (29) and deer reference genomes downloaded from the National Center for Biotechnology Information (NCBI) database (MuD: GCA_020976825; WTD: GCA_023699985). We *de novo* assembled the filtered reads using Megahit v1.2.9 (30) and SPAdes v4.1.0 (metaviral and rnaviral) (31–33). We combined all contigs and deduplicated them (keeping the longest version) using CD-HIT-EST in CD-HIT v4.8.1 (34, 35). We retained contigs larger than 200 bp and performed taxonomical classification by comparing them to the NCBI non-redundant protein database (as of March 2026) using DIAMOND BLASTx v0.9.24 (36), with an e-value cut-off of 1 × 10^−5^. We annotated viral genomes using CenoteTaker3 v3.4.4 (37, 38) followed by manual curation using Geneious Prime® 2026.0.2 (https://www.geneious.com). For downstream analyses, we focused on contigs belonging to viral classes/orders/families or presenting sequence similarity with viruses known to likely infect vertebrate species. We excluded retroviruses (*Retroviridae* family) as it is difficult to determine whether they are endogenous or exogenous.

### Viral abundance calculation

We calculated abundance for each viral taxa as the count of expected reads. We used Bowtie2 v2.4.1 (39) with the “--end-to-end*”* and the “--very-sensitive” options to retrieve the number of trimmed reads mapping to the retained contigs. We calculated overall abundance as the percentage of mapped reads relative to the total number of trimmed reads within the library. We then normalized the abundance by read per million (RPM). When a particular virus was present in several libraries, we applied a threshold relative to the maximum abundance of that virus to reduce the risk of reporting false positives due to index-hopping. Thus, if the total read count in a library was less than 0.1% of the read count in the library where the virus was the most abundant, it was excluded from the analyses (https://www.illumina.com/techniques/sequencing/ngs-library-prep/multiplexing/index-hopping.html).

### Phylogenetic analyses

We aligned amino acid (aa) sequences obtained in this study with representative sequences from viruses of the same genus or family, downloaded from NCBI, using MAFFT v7.490 and default parameters in Geneious Prime (40). We inferred maximum-likelihood phylogenies using IQTREE v2.1.4-beta (41) and ModelFinder to select the best-fitting substitution model (42). We assessed branch support using 1,000 bootstrap replicates using the Ultrafast bootstrap approximation (UFBoot) algorithm (41). For partitioned maximum-likelihood phylogenetic trees (*Orthoherpesviridae*), we generated separated alignments for individual proteins before concatenating them. We visualized phylogenetic trees on Figtree v1.4.4 (http://tree.bio.ed.ac.uk/software/figtree/), then edited them in Inkscape v1.3 (https://inkscape.org). All phylogenetic trees were represented as midpoint rooted for clarity. We applied species demarcation criteria provided by the International Committee on Taxonomy of Viruses (ICTV) to identify putative new viruses (https://ictv.global/report/genome).

### Statistical analyses

We evaluated whether viral richness (number of viral taxa detected) varied among species by using a generalized linear mixed model with a Poisson distribution and a log link function. Species was set as a fixed term, and study site was included as a random effect. We quantified the variance explained by the study site by estimating the random effect variance and by comparing models with and without the random effect using a likelihood ratio test (LRT). We fitted models using the glmmTMB package v1.1.11 in R (43, 44) and evaluated models assumptions and overdispersion using the DHARMa package v0.4.6 (45). We assessed the significance of each variable using the “Anova” function from the car package v3.1.3 (46). We evaluated beta diversity using a Jaccard dissimilarities index, based on the presence/absence of each viral species. We tested differences in viral community composition among deer species and study sites with the “Adonis2” function in the vegan package v2.6-8 (47), using permutational multivariate analyses of variance (PERMANOVA). We verified the distribution of any significant factor using permutational analysis of multivariate dispersions (PERMDISP) using the “betadisper” function. All statistical analyses were conducted in R v4.3.3 (48) and RStudio v2023.6.0.421 (49).

## Results

### Overview of the sequenced libraries

We sequenced a total of 21 pooled RNA libraries built from 210 nasopharyngeal swabs collected from 70 MuD and 140 WTD, across nine states and 14 study sites in the USA during the winter of 2023–2024 (from December to April; **Table 1, Supplementary Table 1**). We generated a total of 992,549,435 read pairs with an average of 47,264,259 read pairs per library (range: 38,337,218–62,353,443 read pairs; **Supplementary Table 2**). After filtering out low- quality reads, rRNA, and host genome, we retained a total of 4,841,984 read pairs with an average of 230,571 per library (range: 27,367–2,951,588). *De novo* assembly (Megahit and SPAdes) produced 228,022 contigs with an average of 10,858 contigs per library (range: 2,646–76,246; **Supplementary Table 2**).

### Viral richness, diversity, and abundance

We identified at least 18 viral classes (**Supplementary Figure 1A**), 28 viral orders (**Supplementary Figure 1B**), and 82 viral families (**Supplementary Figure 1C**). Viruses were detected at every site, but the majority of them were bacteriophages, plant viruses, or protist viruses that we excluded from downstream analyses. When retaining only vertebrate-infecting viruses, we detected lower viral diversity. In total, we identified four viral classes (*Herviviricetes*, *Monjiviricetes*, *Pisoniviricetes* and *Resentoviricetes*), five viral orders (*Herpesvirales*, *Mononegavirales*, *Nidovirales*, *Picornavirales* and *Reovirales*), and seven viral families (*Coronaviridae*, *Orthoherpesviridae*, *Paramyxoviridae*, *Picornaviridae*, *Rhabdoviridae*, *Sedoreoviridae*, and *Tobaniviridae*) (**Figure 1A and 1B, Supplementary Table 3**). We also excluded *Retroviridae* from downstream analyses because of the challenges associated with endogenous retroviruses.

**Figure 1.**
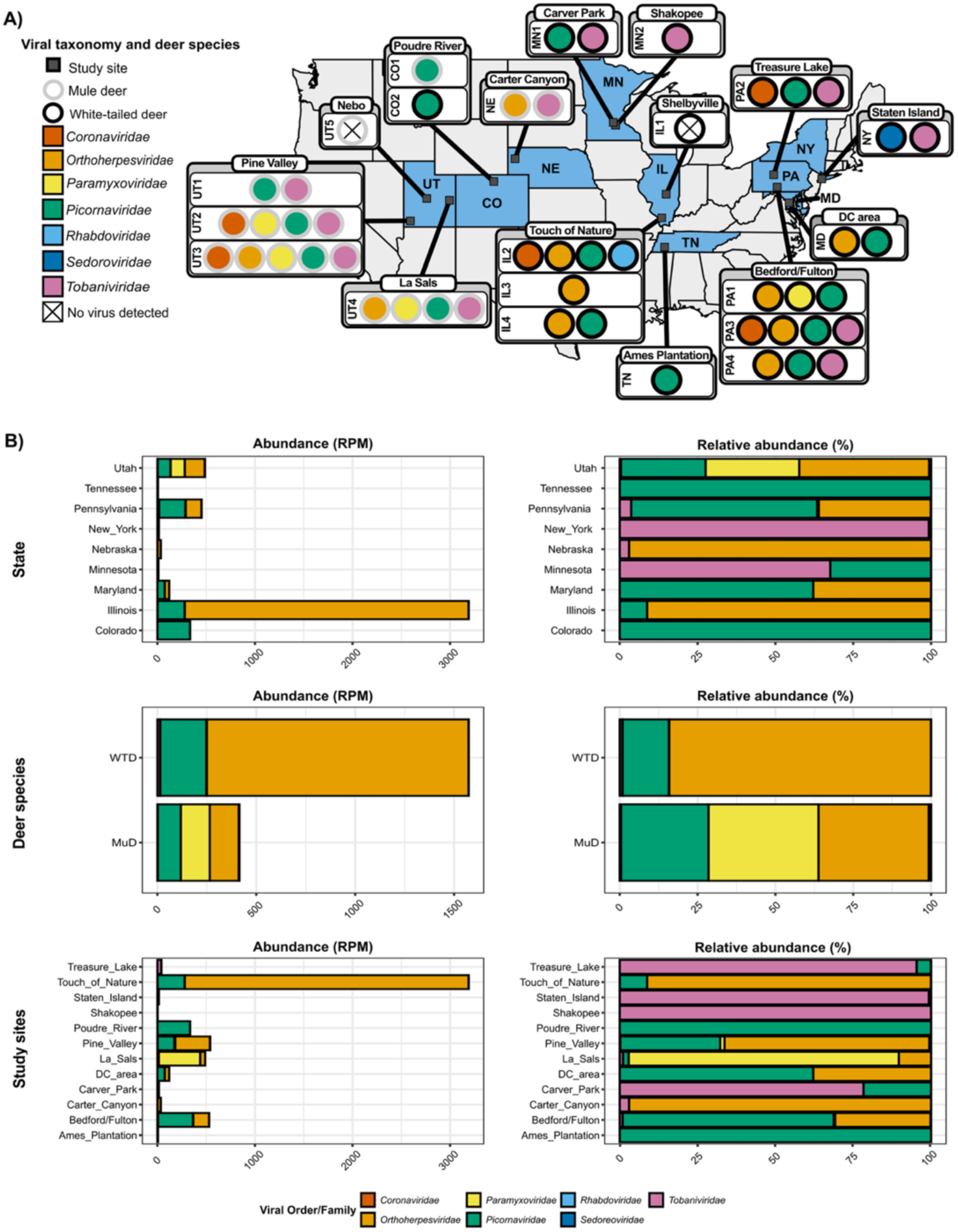
Diversity and abundance of vertebrate-infecting viruses. A) Viral diversity per pool across states and study sites. Study sites are represented by gray squares outlined by a black line. States are abbreviated: CO = Colorado, IL = Illinois, MD = Maryland, MN = Minnesota, NE = Nebraska, NY = New York, PA = Pennsylvania, TN = Tennessee, UT = Utah. For Maryland, DC = District of Columbia. For each study site, colored circles represent identified viral taxa. Circles outlined by a gray line correspond to viruses identified in mule deer, and circles outlined by a black line represent viruses identified in white-tailed deer. Pool names are indicated on the left. B) Abundance per state, deer species and study site. The left plots show overall abundance in reads per million (RPM), and the right plots show relative abundances. For deer species, MuD = mule deer and WTD = white-tailed deer. Only libraries where vertebrate-infecting viruses were found are represented.

These vertebrate-infecting viruses were detected at all sites except Shelbyville, Illinois and Nebo, Utah (**Figure 1A and 1B**). Five of the seven viral families (*Coronaviridae*, *Orthoherpesviridae*, *Paramyxoviridae*, *Picornaviridae* and *Tobaniviridae*) were found across multiple study sites (**Figure 1A and 1B**). Viral richness ranged between zero and five taxa across study sites (**Figure 1A and 1B**). The study site explained negligible variance (*σ^2^* = 0.011) and model comparison showed that including it as a random effect did not improve the model fit (LRT *χ^2^* = 71.818, *df* = 1, *P* = 0.915). Models showed that viral richness was not predicted by the deer species (*χ^2^* = 0.5193, *df* = 1, *P* = 0.4712). Similarly, beta-diversity did not vary among deer species (*pseudo-F* = 0, *R^2^* = 0, *P* = 0). However, viral community composition was different across study sites (*pseudo-F* = 2.18, *R^2^* = 0.76, *P* = 0.023).

### Taxonomic composition of deer virome

#### Coronaviridae

We identified two viruses belonging to the *Coronaviridae* family, *Orthocoronavirinae* subfamily, and *Betacoronavirus* genus. The first one was a bovine-like coronavirus (CoV), that we found in MuD captured in Pine Valley, Utah (**Figure 1A and 1B, Supplementary Table 3**). The virus was detected in two of the three pools available for the study site. We retrieved fragmented genomes consisting of 42 contigs ranging from 232 to 2,482 bp. The most closely related virus was bovine coronavirus BCoV-ENT (AF391541) identified in an individual that suffered fatal pneumonia during a bovine shipping fever epizootic in the USA (50), with which the largest contig, a partial nucleoprotein sequence, shares 99.5% aa identity. Phylogenetic analyses on partial nucleoprotein sequences from members of the *Betacoronavirus* genus, placed the largest sequence obtained in this study in a strongly supported clade, composed of different strains of BCoV, as well as bovine-like CoVs identified in captive wild ruminants (*e.g.*, WTD; sambar deer, *Rusa unicolor*; waterbuck deer, *Kobus ellipsiprymnus*; sable antelope, *Hippotragus niger*; giraffe, *Giraffa camelopardalis*) originating in a wild-animal habitat and a wildlife farm in Ohio, USA (**Figure 2**) (51, 52).

**Figure 2.**
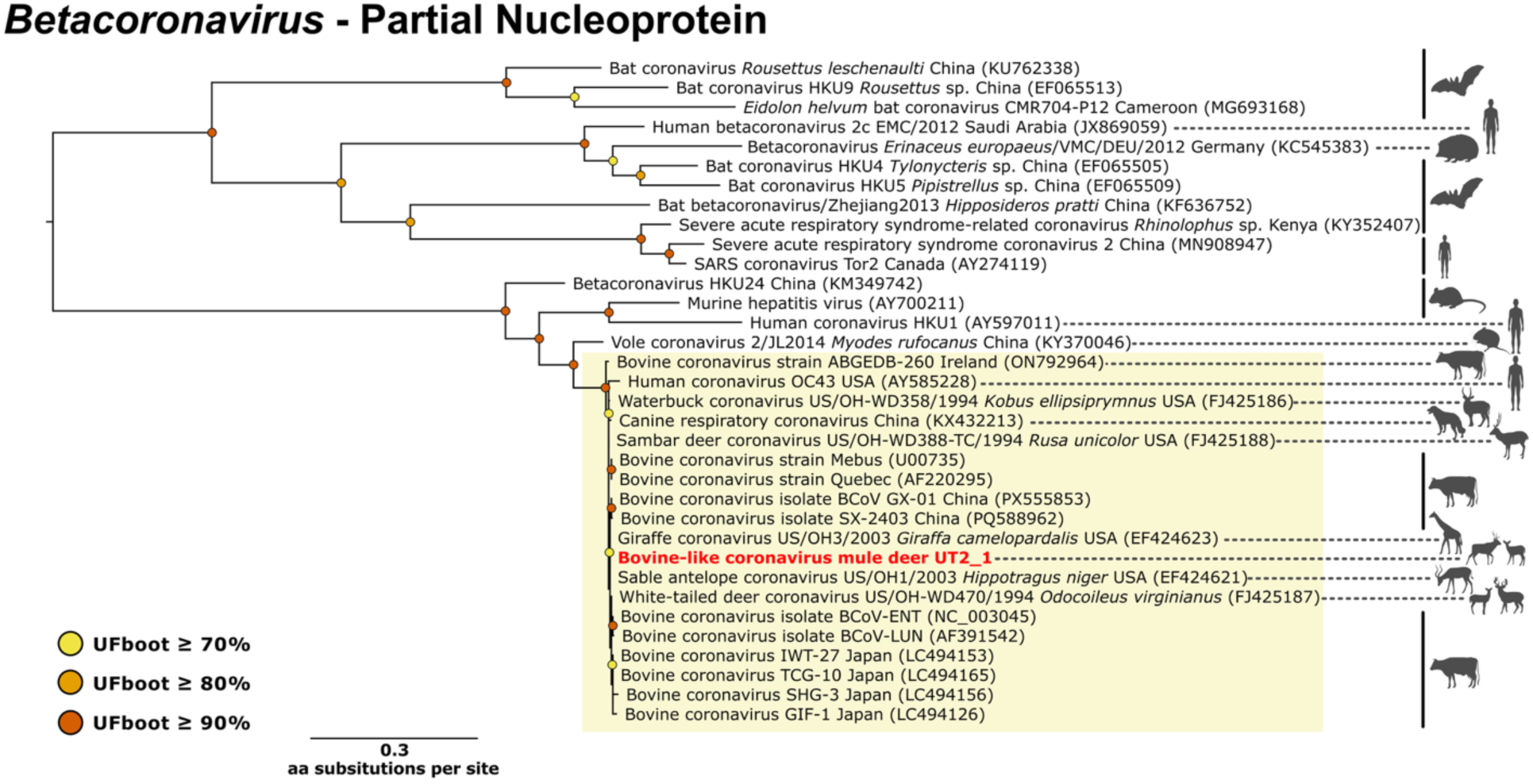
Maximum-likelihood phylogeny of *Betacoronavirus* partial nucleoprotein. The yellow rectangle highlights the clade where the sequence identified in this study clusters. The sequence obtained in this study is highlighted in red.

The second coronavirus we identified was SARS-CoV-2. We detected it in WTD captured in Illinois and Pennsylvania, more precisely in Touch of Nature, Bedford/Fulton and Treasure Lake (**Figure 1A and 1B, Supplementary Table 3**). For Illinois, we identified the virus in one of the three pools available for the study site. We retrieved eleven contigs ranging from 227 to 655 bp that mapped to different regions of the genome. The largest contig was a partial sequence of the nucleocapsid phosphoprotein that shares 100% aa identity with multiple sequences on NCBI (*e.g.*, OM103723, identified in a patient in Mississippi, USA; the sequence is a direct submission to NCBI, and there is no associated reference). For Bedford/Fulton, Pennsylvania, we identified the virus in one of the three pools available and retrieved ten contigs ranging from 228 to 486 bp, also mapping to different regions of the genome. The largest contig was a partial sequence of the ORF1a that shares 100% aa identity with multiple sequences on NCBI (*e.g.*, ON893320, identified in a patient in Texas, USA; the sequence is a direct submission to NCBI, and there is no associated reference). Finally, for Treasure Lake, Pennsylvania, we retrieved two contigs of 241 and 267 bp respectively. They were both partial sequences of the membrane glycoprotein, and the largest contig shares 98.9% aa identity with a sequence isolated from a patient in the USA (PP421552; the sequence is a direct submission to NCBI, and there is no associated reference). For each of the study sites, the contigs were too small and genome coverage too low to identify which SARS-CoV-2 variants were present in the libraries. All of our samples were tested for SARS-CoV-2 by PCR as part of the national-scale disease surveillance network (25). The results of the PCR testing show that we correctly blindly identified SARS-CoV-2 in all sample pools containing positive samples (personal communication).

#### Orthoherpesviridae

We identified members of the *Orthoherpesviridae* family and *Gammaherpesvirinae* subfamily, in multiple libraries (**Figure 1A and 1B, Supplementary Table 3**). We detected them in WTD from Touch of Nature, Illinois (in all pools available), District of Columbia (DC) area, Maryland, and Bedford/Fulton, Pennsylvania (in all pools available). We also detected them in MuD from Carter Canyon, Nebraska, Pine Valley, Utah (in one pool) and La Sals, Utah (**Figure 1A and 1B**). In total, we retrieved 951 contigs ranging from 225 bp to 99,335 bp. The 99,335 bp contig, identified in one of the pools available for Touch of Nature, Illinois, was a near-complete genome that covered the complete unique long (UL) region. Using CenoteTaker3, we were able to reconstitute another near-complete genome for another pool originating from the same study site. The sequence is 114,549 bp in length and contained the complete UL region along with a portion of the unique short (US) region. We also retrieved a partial genome of 56,144 bp, containing a portion of the UL region for Pine Valley, Utah. Partitioned phylogeny inferred from six conserved genes for members of the *Gammaherpesvirinae* subfamily (53), showed that the sequences clustered together within a highly supported clade containing viruses of the *Macavirus* genus, identified in other ruminants (**Figure 3**). When performing BLAST searches, the sequences also showed high degree of similarity with other unclassified herpesviruses for which only partial sequences were available. One of them was type 2 ruminant *Rhadinovirus* of mule deer, identified in MuD and WTD in Canada (sequences were deposited directly in NCBI and we found no associated publications). The largest sequence available was 3,690 bp long (HM014314) and comprised partial coding sequences for the glycoprotein B, and the DNA polymerase, with which our sequences share 99.53% and 98.40% aa identity, respectively.

**Figure 3.**
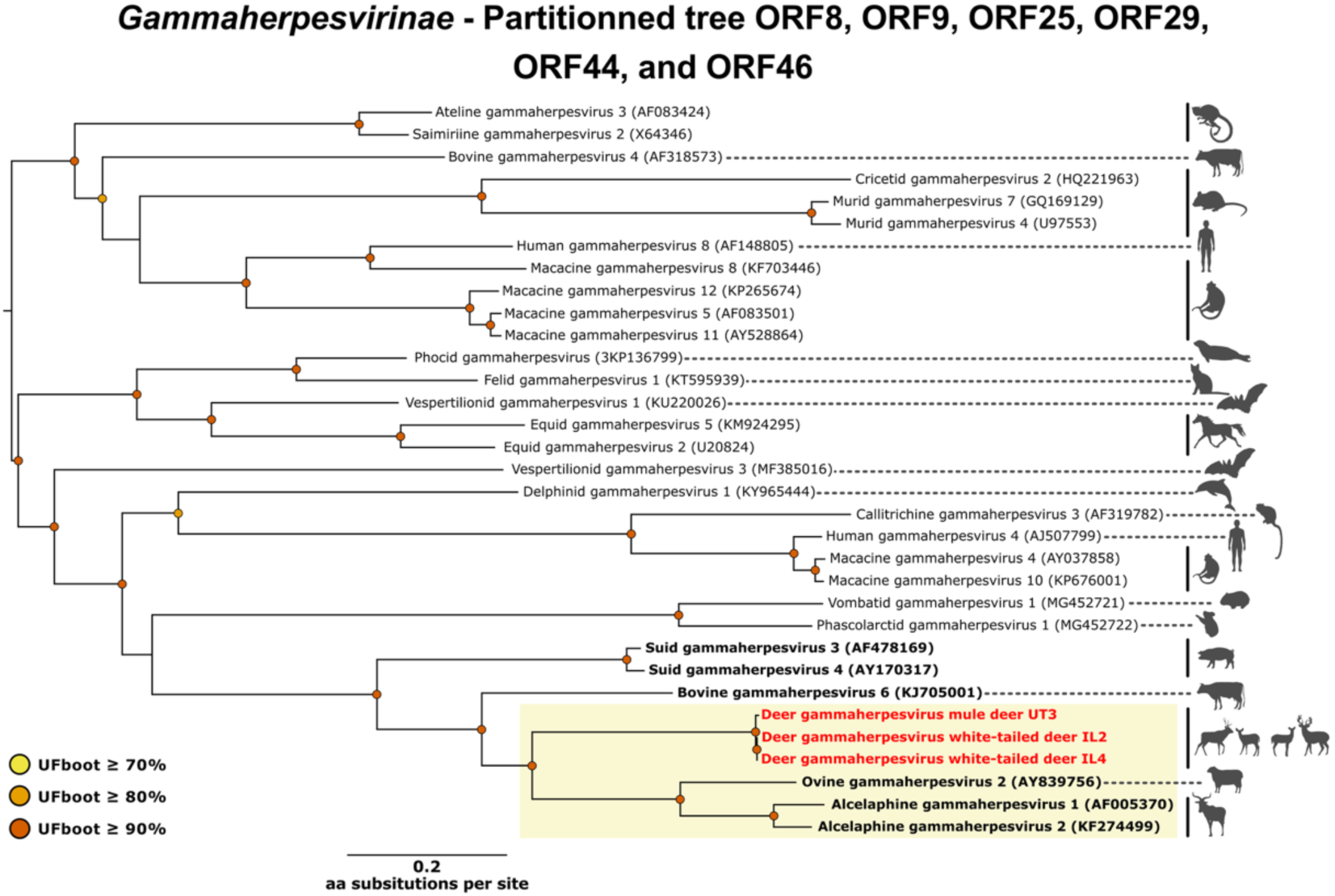
Partitioned maximum-likelihood phylogeny of *Gammaherpesvirinae* concatenated ORF8, ORF9, ORF25, ORF29, ORF44 and ORF46 amino acid alignments. The yellow rectangle highlights the clade where sequences identified in this study cluster. Sequences obtained in this study are highlighted in red. Sequences belonging to the *Macavirus* genus are highlighted in bold. ORF8 encodes for envelope glycoprotein B, ORF9 for DNA polymerase catalytic subunit, ORF25 for major capsid protein, ORF29 for DNA packaging terminase subunit 1, ORF44 for helicase-primase helicase subunit, and ORF46 for uracil-DNA glycosylase.

#### Paramyxoviridae

We identified a virus belonging to the *Paramyxoviridae* family, *Orthoparamyxovirinae* subfamily and *Respirovirus* genus. We identified it in MuD from Pine Valley, Utah (in two pools), La Sals, Utah, and WTD from Bedford/Fulton, Pennsylvania (in one pool) (**Figure 1A and 1B, Supplementary Table 3**). The virus was closely related to bovine parainfluenza virus 3 (BPIV-3).

For Pine Valley and La Sals, Utah, we identified near-complete genomes measuring 15,409 and 15,465 bp, respectively. For Pennsylvania, we only retrieved genome fragments ranging from 232 bp to 589 bp. The near-complete sequences share 93.7 and 97.2% aa identity with the closest BPIV-3 published sequence (AF178654) (54). Phylogenetic analyses of the large (L) protein, used by the ICTV for species demarcation, placed the two sequences obtained in this study together within a clade comprising multiple BPIV-3 sequences. This clade was sister to a second clade containing BPIV-3 and human respirovirus 3 sequences (**Figure 4**). The sequences obtained in this study share 92.1 and 92.3% aa identity on the L protein with the other unique published BPIV-3 like sequence isolated from a deer species (ON014594) (55) (**Figure 4**). The virus was detected in a fallow deer fawn (*Dama dama*) that died during a mortality event involving three other fawns and three adult hinds in a deer herd in England, United Kingdom (55). Although it was not possible to establish a link between the infection and transmission from cattle, the deer fawns shared a shed with young cattle that were reported to be coughing (55).

**Figure 4.**
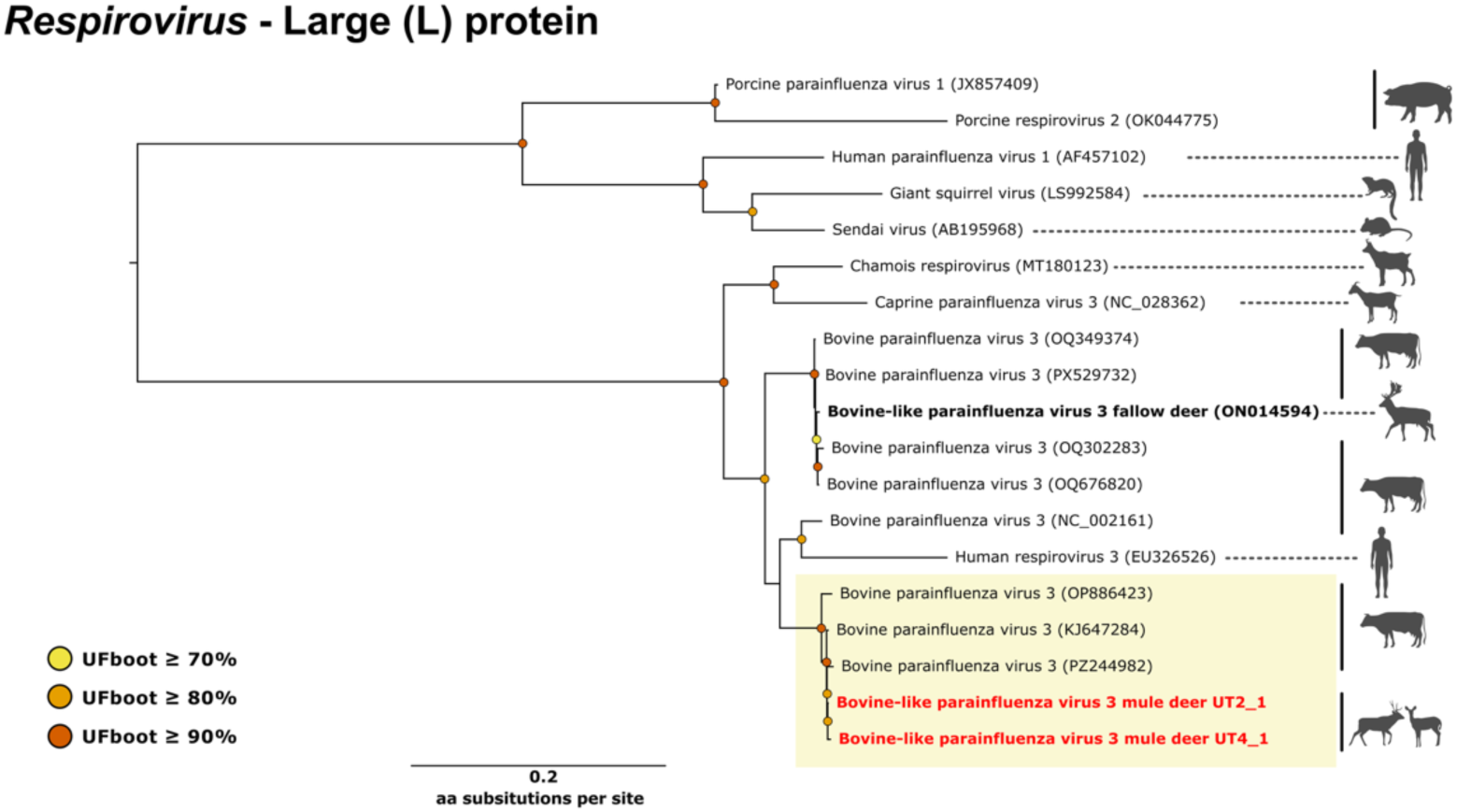
Maximum-likelihood phylogeny of *Respirovirus* large (L) protein. The yellow rectangle highlights the clade where sequences identified in this study cluster. Sequences obtained in this study are highlighted in red. The only other published sequence of bovine parainfluenza virus 3 detected in a deer species is highlighted in bold.

#### Picornaviridae

We identified a virus belonging to the *Picornaviridae* family, *Caphthovirinae* subfamily and *Aphthovirus* genus in both species, across seven states and nine study sites (**Figure 1A and 1B, Supplementary Table 3**). We detected the virus in MuD from Poudre River, Colorado, Pine Valley, Utah (all pools), and La Sals, Utah. We also identified it in WTD from Poudre River, Colorado, Touch of Nature, Illinois (two pools), DC area, Maryland, Carver Park, Minnesota, Bedford/Fulton, Pennsylvania (all pools), Treasure Lake, Pennsylvania, and Ames Plantation, Tennessee (**Figure 1A and 1B**). In total, we recovered 13 complete coding sequences. For some of the pools, we recovered multiple distinct near-complete genomes (**Figure 5**). Bovine rhinitis A virus (BRAV) was the closest relative. The 13 sequences share between 67.2% and 68.5% aa identity with the polyprotein of multiple published BRAV sequences (*e.g.*, KU159364, KP264974, KT948520) (56–58). Phylogenetic analyses of the polyprotein and the RNA-dependent RNA polymerase aa sequences from members of the *Aphthovirus* genus revealed that all sequences identified in this study clustered in a distinct clade, sister to the clade containing all BRAV sequences (**Figure 5A and 5B**). This monophyletic clade, combined with the aa similarity, suggests that these represent a putative new deer derived virus clade (59). The deer clade consisted of two subclades (**Figure 5A and 5B**). The first one contained only sequences isolated from WTD, whereas the second contained all sequences isolated from MuD and one sequence isolated from WTD captured in Poudre River, Colorado, where both species live in sympatry.

**Figure 5.**
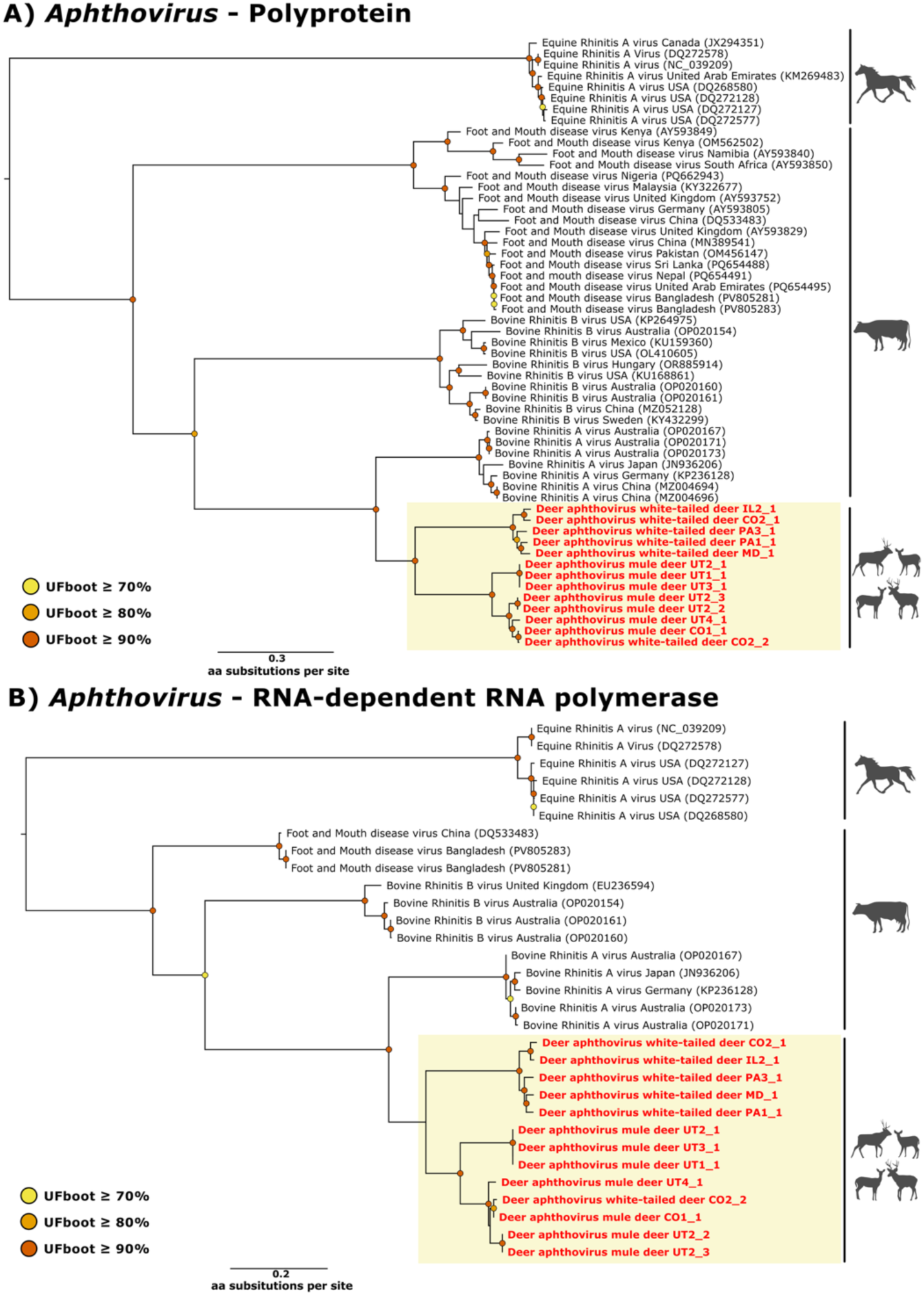
Maximum-likelihood phylogeny of *Aphthovirus* A) Polyprotein and B) RNA-dependent RNA polymerase protein. Yellow rectangles highlight the clade where sequences identified in this study cluster. Sequences obtained in this study are highlighted in red.

#### Rhabdoviridae

We identified one virus belonging to the *Rhabdoviridae* family, *Alpharhabdovirinae* subfamily, and *Ephemerovirus* genus. The virus was detected in WTD from Touch of Nature, Illinois (in one pool) (**Figure 1A and 1B, Supplementary Table 3**). We recovered six contigs ranging from 238 to 968 bp. The closest match was New Kent County virus (NKCV; MF615270) identified in blacklegged ticks (*Ixodes scapularis*) collected in Virginia, USA (60). The largest contig shares 89.37% aa identity with NKCV on a partial sequence of the nucleoprotein. Phylogenetic analyses of partial nucleoprotein of members of the *Ephemerovirus* genus (**Figure 6**) confirmed that the sequence obtained in this study clusters with NKCV in a clade sister to Koolpinyah virus (KM085029) identified in Australian cattle (61), and Kotonkan virus (HM474855) identified in *Culicoides* spp. biting midges in Nigeria (62). However, robust phylogenetic analysis could not be undertaken due to short contig size and limited genetic information available for this group.

**Figure 6.**
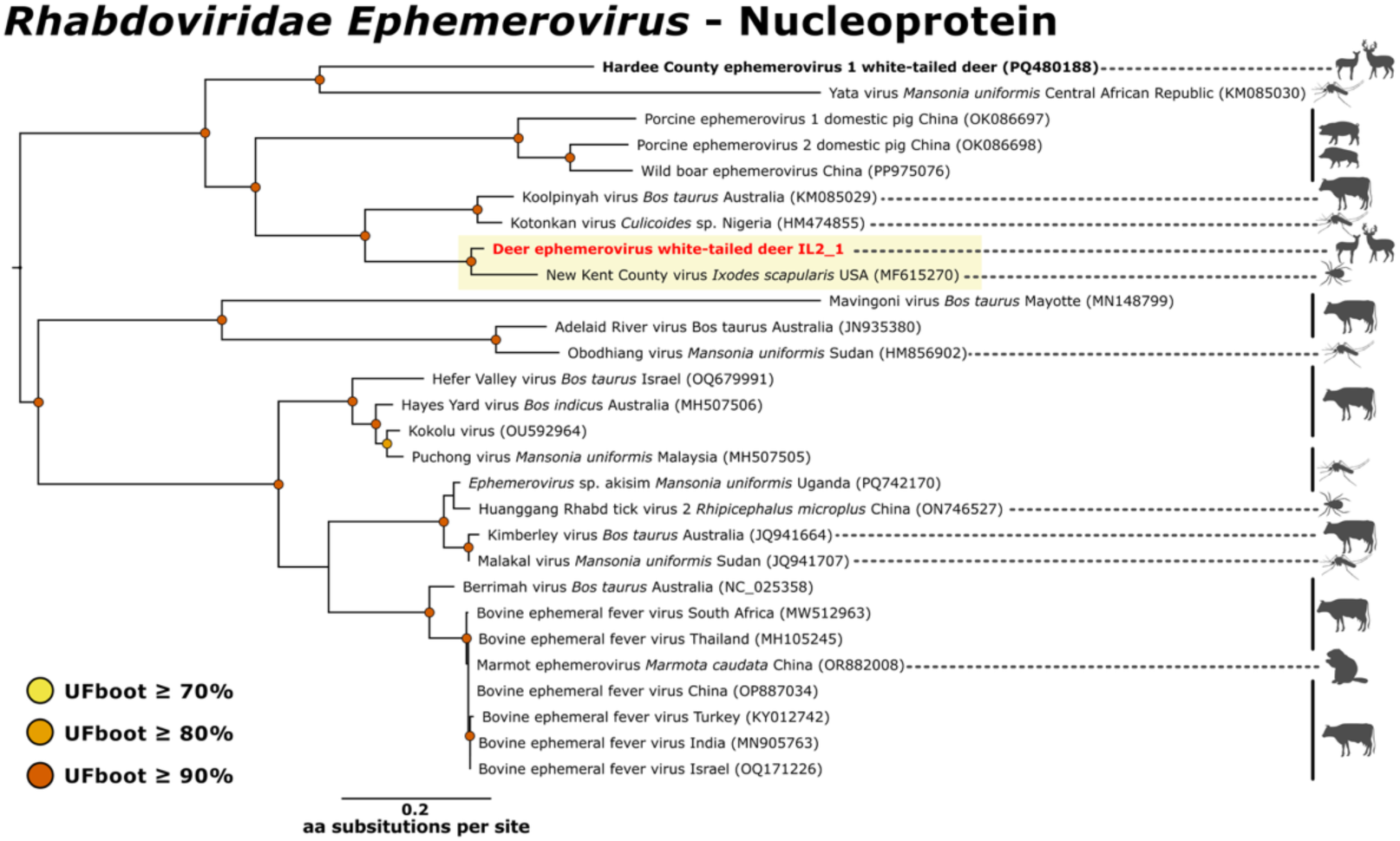
Maximum-likelihood phylogeny of *Ephemerovirus* partial nucleoprotein. The yellow rectangle highlights the clade where the sequence identified in this study clusters. The sequence obtained in this study is highlighted in red. Hardee County ephemerovirus 1, only other ephemerovirus identified in a deer species using molecular tools is highlighted in bold.

#### Sedoreoviridae

We identified one virus belonging to the *Sedoreoviridae* family and *Orbivirus* genus in WTD from Staten Island, New York (**Figure 1A and 1B, Supplementary Table 3**). We retrieved two contigs of 258 and 296 bp, which share 100% aa identity, respectively, with segment 5 and segment 1 of CHeRI orbivirus 1 (MK903626) identified in WTD in the USA (63). To further explore the presence of this virus we mapped trimmed reads from the New York library against complete sequences of other CHeRI orbivirus 1 segments (63), and retrieved additional reads mapping to segment 1 (MK903619, six reads), segment 2 (MK903621, two reads), segment 4 (MK903622, two reads), and segment 5 (MK903626, ten reads), supporting the presence of the virus within the library.

#### Tobaniviridae

We identified a member of the *Tobaniviridae* family in multiple libraries (**Figure 1A and 1B, Supplementary Table 3**). We detected the virus in MuD captured in Carter Canyon, Nebraska Pine Valley, Utah (all pools) and La Sals, Utah. We also detected it in WTD from Carver Park, Minnesota, Shakopee, Minnesota, Staten Island, New York, Bedford/Fulton, Pennsylvania (two pools) and Treasure Lake, Pennsylvania (**Figure 1A and 1B**). In total, we retrieved 172 contigs ranging from 225 to 8,939 bp. Using CenoteTaker3 we were able to reconstitute a 20,466 bp near-complete genome for Treasure Lake, Pennsylvania. The most closely related viruses were bovine nidovirus 1 (BoNV; ON330451) (64) and ovine nidovirus (OR885915) (65), with which the sequence shares between 61.74 and 62.34% identity, suggesting a putative new species. Phylogenetic analyses of the conserved polyprotein 1b from members of the *Tobaniviridae* family placed the sequence identified in this study near a strongly supported clade constituted of BoNV and ovine nidovirus sequences (**Figure 7**).

**Figure 7.**
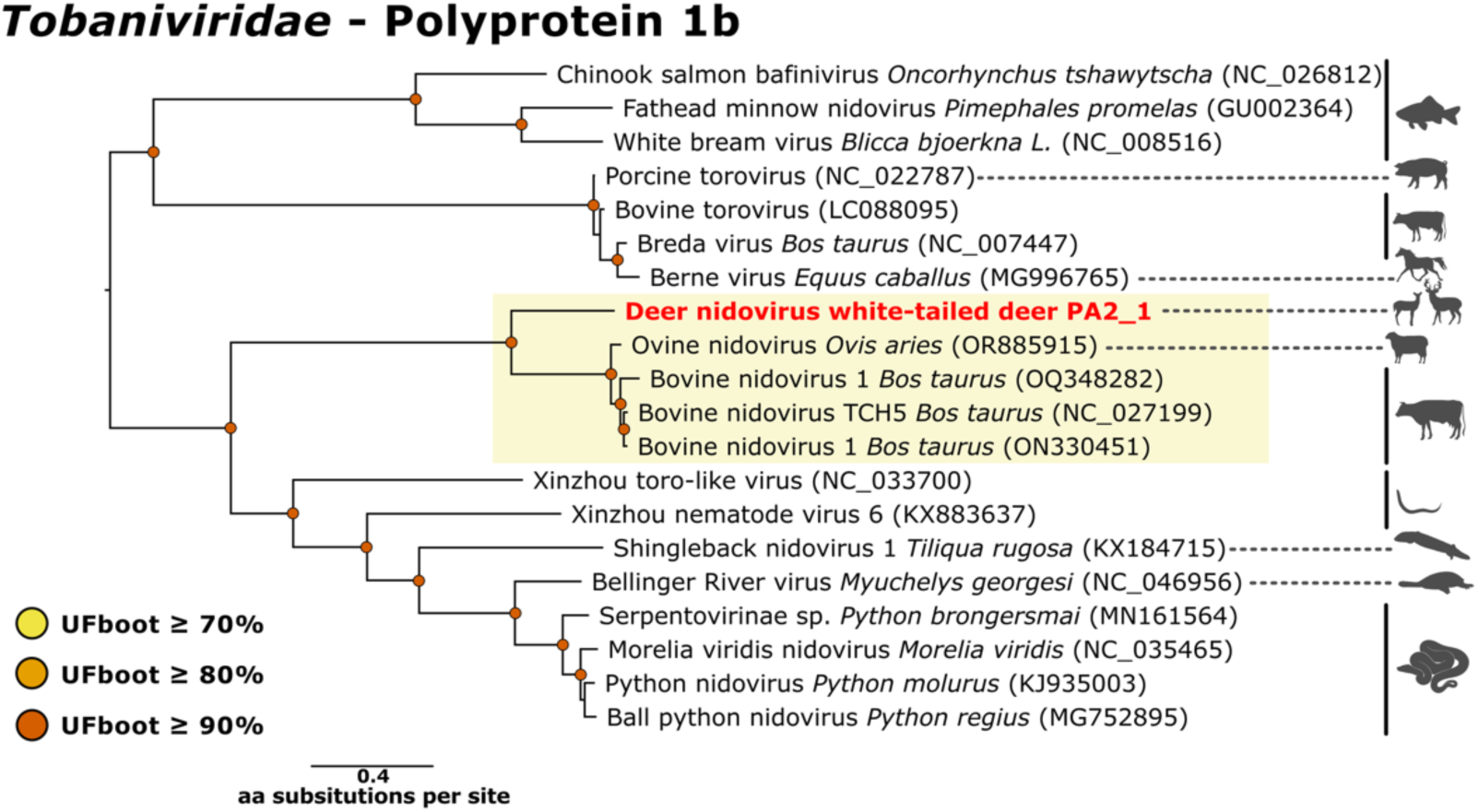
Maximum-likelihood phylogeny of *Tobaniviridae* Polyprotein 1b. The yellow rectangle highlights the clade where the sequence identified in this study clusters. The sequence obtained in this study is highlighted in red.

## Discussion

Our metagenomic high-throughput sequencing approach shed light on the upper respiratory virome of live MuD and WTD, captured across the USA. We identified known and novel viruses, including a novel *Aphthovirus* (*Picornaviridae* family) and nidovirus (*Tobaniviridae* family), for which we recovered near-complete genomes. We also documented several viruses that originate in, or are closely related to, viruses previously described in humans or livestock, supporting viral spillover at the deer–human–livestock interface.

The novel deer *Aphthovirus* was found in both MuD and WTD across multiple states and study sites, suggesting that this previously undescribed virus may circulate widely in deer populations in the USA. Phylogenetic analyses revealed clustering by deer species, indicating possible host-adapted strains or variants. However, we also documented a potential cross-species transmission. One of the sequences originating from the WTD pool of Poudre River, Colorado, clustered with a sequence originating from the MuD pool of the same study site, within the subclade dominated by MuD sequences. Host specificity and transmission mechanisms for this virus remain unknown at this time. However, field notes indicate that one of the WTD, from which a sample was included in the pool for Poudre River, had been observed with the MuD herd for more than two weeks before capture. Individual-level molecular surveys, full genome sequencing, and phylodynamic studies could lead to a better understanding of the ecology, epidemiology, and evolutionary history of this new virus (66). The genus *Aphthovirus* encompasses multiple viruses of health significance for domestic animals, like BRAV, or foot-and-mouth disease virus (56–58). These data could prove valuable in inferring models to predict the transmission of these other viruses (67).

Bovine-like CoVs are commonly detected in wild ruminants, including deer. They share strong biological, genetic, and antigenic similarities with BCoV (51, 68–70). However, reliable genetic markers to distinguish them from BCoV are currently lacking (51, 68). Previous studies have suggested ongoing viral exchange and possible co-evolution between livestock and wild ruminants, with wild ruminants potentially serving as natural reservoirs (68). Here, a bovine-like CoV was identified in deer captured in Pine Valley, Utah, where cattle graze freely on private and public lands, providing opportunities for contacts between deer and cattle. Although the virus we detected shared a strong identity with cattle viruses, we lack sufficient data to determine its origins. We also identified a bovine-like PIV-3 in deer captured in Pine Valley, Utah, La Sals, Utah, and Bedford/Fulton, Pennsylvania. Although free-range cattle farming is less common in Pennsylvania than Utah, Bedford and Fulton counties are home to many farms and ranches where cattle are raised on pasture. Bovine-like PIV-3 is commonly reported in domestic and wild ungulates. In the USA, serological evidence of the virus has for example been detected in MuD (71), WTD (72, 73), or bighorn sheep (*Ovis canadensis*) (74, 75). In England, United Kingdom, a study reported the molecular detection of a bovine-like PIV-3 during a mortality event in a fallow deer herd (55). However, no clear association has been established between bovine-like PIV-3 and diseases in deer. Bovine-like CoVs have been reported in healthy or diarrheic wild ruminants. They have not been associated with respiratory disease outbreaks in these species (68). In contrast, BCoV and BPIV-3 are recognized causes of serious diseases in cattle, including the bovine respiratory disease complex (76–79). Given close phylogenetic relationships between deer and cattle, a clearer understanding of transmission dynamics and the potential risks of spillback from deer to livestock is warranted.

Several herpesviruses of the genus *Macavirus* are responsible for fatal illnesses in ruminants (*e.g.*, malignant catarrhal fever) (56–59). Although the virus detected here likely belongs to this genus, it was also nearly identical to type 2 ruminant *Rhadinovirus* of mule deer, suggesting that it may be the same virus. The limited genetic data available on type 2 ruminant *Rhadinovirus* of mule deer may have contributed to an incorrect taxonomic classification. Herpesviruses are large and complex DNA viruses challenging both their discovery and taxonomical classification. Interestingly, it was surprising to retrieve near-complete herpes genomes in our dataset given our RNA-sequencing approach. These viruses were relatively abundant in our libraries. Because we applied a mild DNase treatment to our samples (27), it is likely that we did not remove all DNA molecules. Alternatively, during the lytic phase (active replication), gammaherpesviruses are known to undergo extensive genome-wide transcription (84, 85). We may have been able to reconstitute near-complete genomes thanks to overlapping transcripts. The use of DNA sequencing techniques could enable the sequencing of the missing regions for this virus and shed light on other DNA viruses present in the libraries, especially during their latent stages.

The putative new nidovirus identified in both species, shared similarities with BoNV and ovine nidovirus, reported in cattle (*e.g.*, USA, Canada, Australia) or sheep (Hungary) presenting respiratory symptoms (4, 64, 65, 86, 87). Discovered thanks to metagenomics, BoNV and ovine nidovirus remain relatively poorly understood. Often found in coinfection with other respiratory pathogens, they have not been directly linked to respiratory diseases (65, 86, 87). To the best of our knowledge, this is the first report of a member of the *Tobaniviridae* family in a deer species. These findings emphasize the necessity to elucidate their ecology and pathogenicity. Nidoviruses (*Nidovirales* order) and especially some members of the *Tobaniviridae* family (*e.g.*, bovine torovirus, porcine torovirus) are known to perform recombination (88–90), and potentially cross species barriers (90, 91). Given the circulation of several nidoviruses among phylogenetically related species, these data are essential to understand the risk of emergence of novel viruses that could be responsible for serious diseases in domestic animals and wild deer populations.

The *Ephemerovirus* detected in Touch of Nature, Illinois, was related to NKCV, discovered in blacklegged ticks collected in Virginia, USA (60). *Ephemerovirus* are arboviruses usually transmitted by mosquitoes or biting midges and primarily infecting ruminants (92). Bovine ephemeral fever virus, pathogenic in cattle (93), has been previously reported in deer based on serology (*e.g.*, 54). However, Hardee County ephemerovirus 1 is the only *Ephemerovirus* reported in cervids using molecular tools (95). The virus was discovered in a deceased farmed WTD in Florida, USA, but the mortality could not directly be attributed to the virus (95). CHeRi orbivirus 1 found in New York deer, also belongs to a viral genus composed of arboviruses. The virus is one of the four CHeRI orbiviruses recently discovered in dead farmed WTD in Florida, and Pennsylvania, USA (95, 63, 96). Several viruses among the *Orbivirus* genus can cause severe diseases in deer, including epizootic hemorrhagic disease virus (EHDV) and bluetongue virus (97); however, the pathogenicity of these CHeRI orbiviruses is yet to be determined. Although investigators have identified them in dead animals, they have only been found in coinfection with other viruses, such as EHDV or Hardee County ephemerovirus 1 (95, 63).

The deer species was not a predictor for viral richness or beta diversity. However, the composition of viral communities changed across study sites as shown in other studies (98–102), suggesting that environmental factors may be the main driver for viral distribution and composition in deer across the USA. Seasonality plays a crucial role in infection dynamics (103) and therefore in shaping viral communities (104). Here, we focused on samples collected during winter. It would be valuable to compare viral communities over time to identify inter-season or inter-annual variation. While we investigated viral communities of deer upper respiratory tracts, testing for other sample types, easily collected during captures (*e.g.*, saliva, feces), may provide valuable data on viruses present in other niches. Only one of our study sites included samples collected from sympatric deer species. It would be beneficial to include more study sites where multiple deer species co-exist. It would also be relevant to explore the virome of other deer species across the USA (*e.g.*, elk, *Cervus canadensis*, moose, *Alces alces*) as they may present different ecology and therefore differences in their viral communities.

This study represents the first large-scale characterization of the upper respiratory virome of free-ranging mule deer and white-tailed deer across the USA. By identifying both previously recognized viruses and several putative new viruses, our findings expand current knowledge of viral diversity in wild cervids and establish an important baseline for future surveillance. These data provide a foundation for developing targeted molecular diagnostics to measure prevalence as well as study the epidemiology and evolutionary factors that shape viral communities in deer. The detection of viruses closely related to those circulating in livestock, together with evidence of widespread novel deer viruses, highlights the potential to further elucidate viral dynamics at the deer–human–livestock interface. As deer populations continue to overlap with agricultural landscapes and human communities, understanding the diversity, transmission dynamics, and evolution of their viruses will be essential for anticipating pathogen emergence, improving wildlife health, and the development of management interventions to reduce disease risks across species.

## Supporting information

Supplementary Table 1: Details on samples included in the different libraries/pools.

Supplementary Table 2: Details on reads and contigs produced for each library/pool.

Supplementary Table 3: Details on number of viral contigs, length, and closest viral match identified for each library/pool.

Supplementary Figure 1. General taxonomic composition by A) Viral Classes, B) Viral orders, and C) Viral families.

## Acknowledgements

We thank Cara Brennan, Casey Maynard, Sarah Way, and Michelle Gibison with the Wildlife Futures Program for their help with sample receipt, management, and shipment logistics. We thank Lisa Mattei and Daniel Cutillo from the Center for Host-Microbial Interactions, respectively for their help in troubleshooting library preparation and for sequencing the libraries. We also thank those not listed among the authors who contributed to the extensive field effort and sample acquisition. The findings and conclusions in this manuscript are those of the authors and should not be construed to represent any official USDA or United States (U.S.) Government determination or policy. Any use of trade, firm, or product names is for descriptive purposes only and does not imply endorsement by the U.S. Government.

## Data availability

Raw reads (fastq files) generated in this study are available in the NCBI Short Read Archive (SRA) database under BioProject PRJNA1477166 (Biosamples accession numbers: SAMN60746848–SAMN60746868). Assembled viral sequences were deposited in GenBank under accession numbers PZ564555–PZ564596, and PZ649434–PZ649458.

## Ethics statement

Samples used in this study were obtained as part of a national longitudinal study conducted throughout the United States of America. Within each state, each research team established separate but similar Institutional Animal Care and Use Committee protocols approved by their academic institutions (Colorado State University: #3872 and #5002 for Northern Colorado and Nebraska sites, respectively; Southern Illinois University: #21–028; University of Maryland: #FR-AUG-23-37; University of Minnesota: #2209-40440A; Columbia University: #AC-AABT8667; Pennsylvania State University: #202202225; University of Tennessee Knoxville: #2850–1021; Utah State University: #13045).

## Supplementary Tables and Figures

### Tables

**Supplementary Table 1: Details on samples included in the different libraries/pools.**

**Supplementary Table 2: Details on reads and contigs produced for each library/pool.**

**Supplementary Table 3: Details on number of viral contigs, length, and closest viral match identified for each library/pool.**

### Figures

**Supplementary Figure 1.**
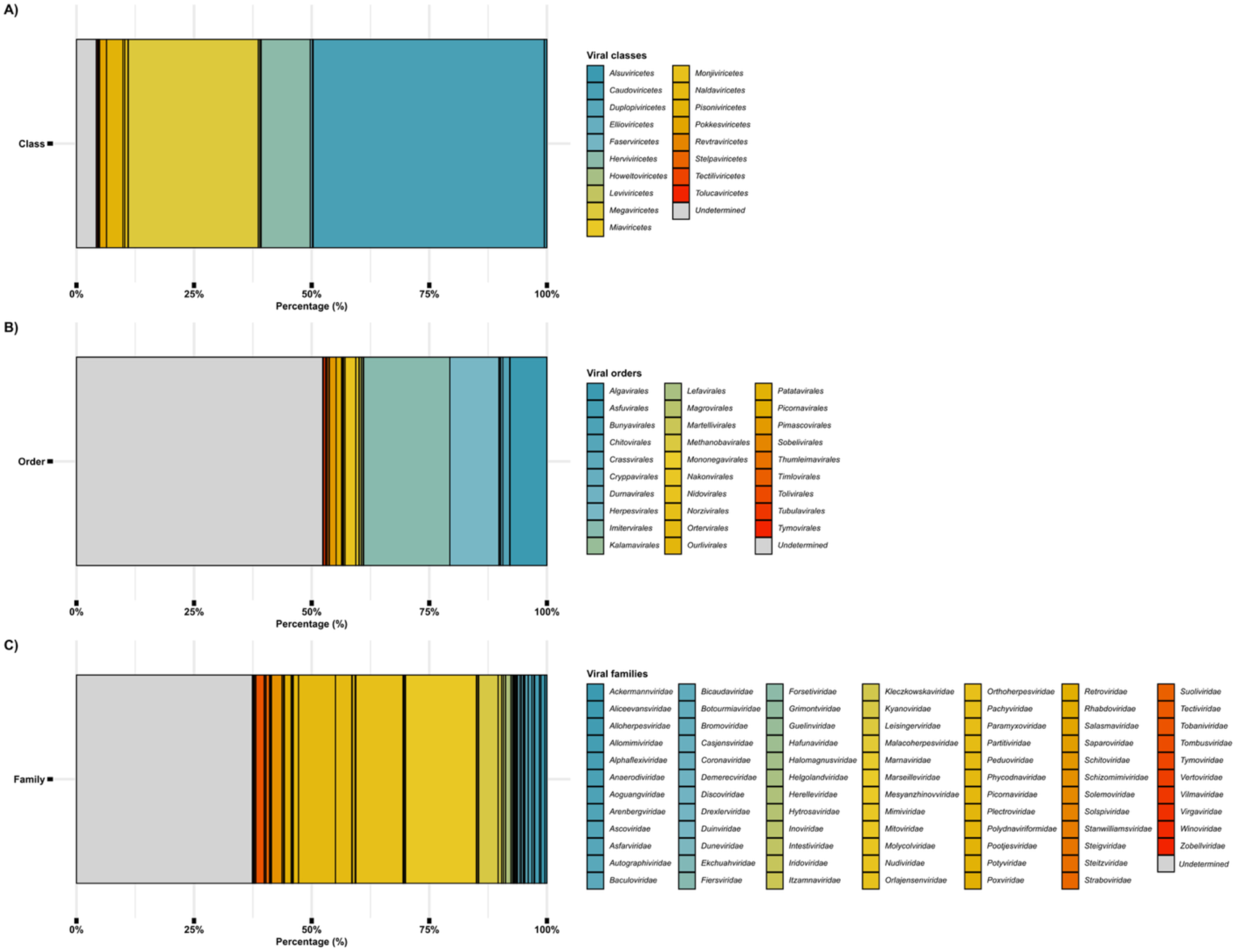
General taxonomic composition by A) Viral Classes, B) Viral orders, and C) Viral families. The figure represents the percentage of contigs assigned to each viral operational taxonomic unit (vOTU) in relation to all the contigs for which DIAMOND BLASTx identified a viral hit. “Undetermined” encompasses viruses for which higher taxonomic rank (*e.g.*, Realm, Kingdom, Phylum) may be available but not the taxonomic rank represented on the figure, or viruses that the International Committee on Taxonomy of Viruses (ICTV) has not officially assigned to a specific vOTU.

