## Supplementary figures and images for "Widespread occurrence of bovine-like and new viruses in wild deer across the United States"

### Supplementary Figure 1. General taxonomic composition by A) Viral Classes, B) Viral orders, and C) Viral families.

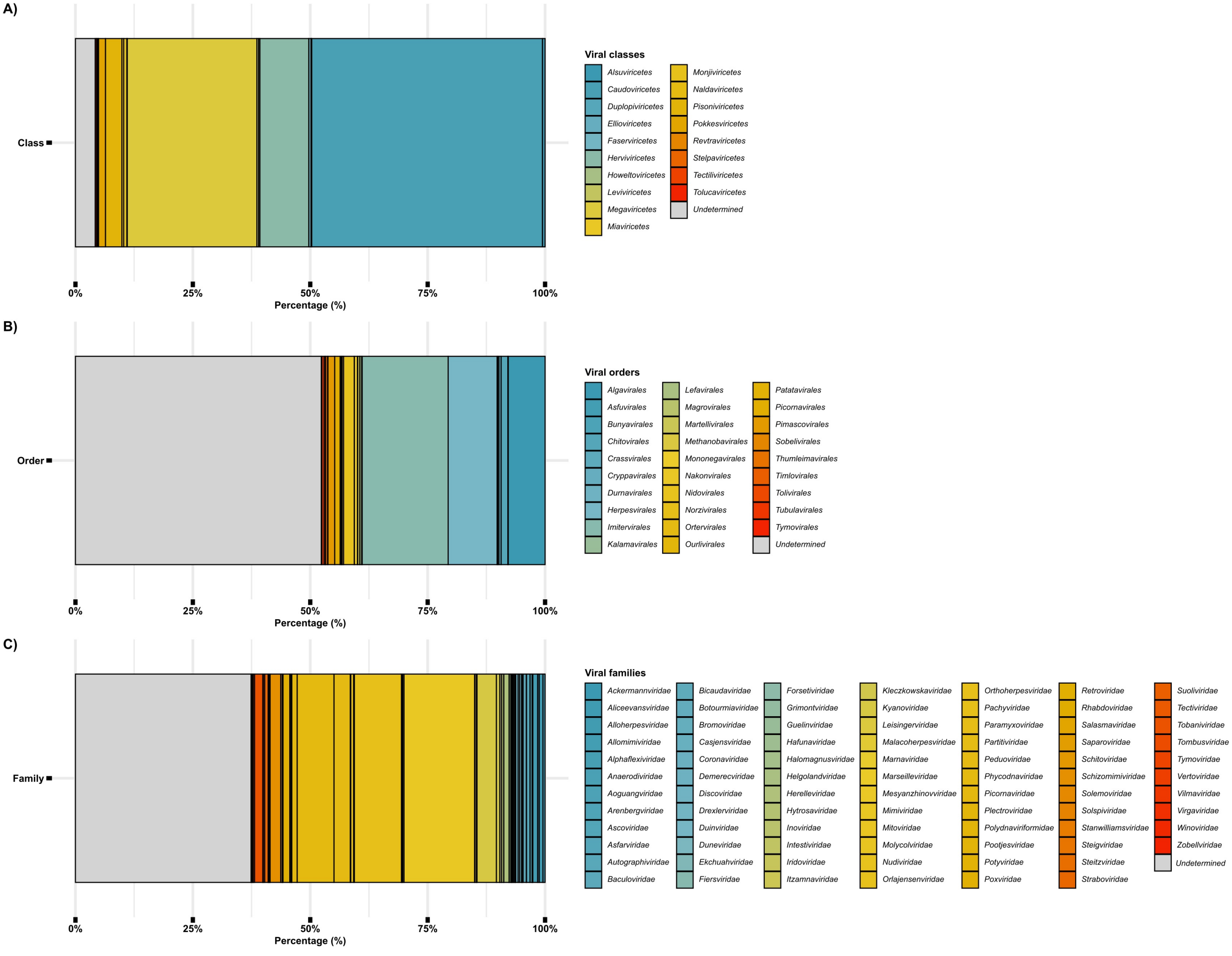
